# Testing the pathogenic potential of *Cryphonectria parasitica* and related species on three common European Fagaceae

**DOI:** 10.1101/2020.03.17.995100

**Authors:** Francesca Dennert, Daniel Rigling, Joana B. Meyer, Christopher Schefer, Eva Augustiny, Simone Prospero

## Abstract

Invasions by non-native pathogens represent a major threat to managed and natural ecosystems worldwide. Although necessary for adopting preventive strategies, the identification of invasive species before they are introduced is particularly difficult. Indeed, most pathogenic species that have become established in the last decades were first described only after they became invasive. To prevent further biological invasions, not only the early identification of potential new invasive plant pathogens is crucial, but also the assessment of their potential host range. In this study, we determined the pathogenicity and the saprotrophic ability of three *Cryphonectria* species towards three potential hosts in the family Fagaceae. For this, seedlings and dormant stems of European chestnut (*Castanea sativa*), pedunculate oak (*Quercus robur*) and European beech (*Fagus sylvatica*) were inoculated with different genotypes of *C. parasitica* (Asian species, invasive in Europe), *C. naterciae* (European species), and *C. japonica* (Asian species, not present in Europe). Lesion growth was measured and mortality assessed for four months. The highest damage was caused by *C. parasitica* on European chestnut, while *C. japonica* and *C. naterciae* induced significantly smaller lesions on this host species. All three *Cryphonectria* species did not grow saprophytically on *F. sylvatica* and *Q. robur*, but successfully colonized dormant stems of *C. sativa*. In the context of biological invasions, our study shows that the Asian *C. japonica* most likely represents a much less severe threat than *C. parasitica* for the tested European host species. Nonetheless, the ability of *C. naterciae* and *C. japonica* to saprotrophically colonize fresh chestnut wood may suggest that they could become established in chestnut forests and eventually infect weakened chestnut trees or other hosts not tested in this study.

## 1 Introduction

The risk of accidentally moving plant pathogens outside their native range is constantly increasing because of global trade and other anthropogenic activities (Santini et al., 2013). Climatic changes, which allow pests to survive in formerly climatic adverse regions, further contribute to range expansions (Stenlid and Oliva, 2016). In the new range, harmful organisms frequently encounter new plant species, some of which may be highly susceptible. In this case, the introduced pest can become invasive (Ghelardini et al., 2017). Detecting invasive species before they are introduced is particularly difficult. Indeed, most pathogenic species that have become established in the last decades were first described only after they became invasive (Brasier, 2008). Assessing the pathogenicity of related species of already present invasive pathogens may be a valuable strategy to spot potentially invasive species before they are introduced (Gilbert et al., 2015).

Although plant pathogens in the introduced range generally attack species which are closely related to the primary host in the natural range (Gilbert and Webb, 2007; De Vienne et al., 2009; Gilbert et al., 2015), in some cases drastic host changes are observed. For example, the rust fungus *Puccinia psidii* considerably expanded its host range when introduced to Australia (Carnegie and Lidbetter, 2012). The bark pathogen *Chrysoporthe cubensis* causes severe damage on introduced *Eucalyptus* spp. and native Myrtaceae and Melostomataceae in Southeast Asia, South America, Africa and Australia (Rodas et al., 2005; Nakabonge et al., 2007; Pegg et al., 2010). Therefore, to prevent further biological invasions not only the early detection of potential new invasive plant pathogens is crucial, but also the assessment of its potential host range.

*Cryphonectria parasitica*, the causal agent of chestnut blight, is one of the most damaging invasive pathogens in forest ecosystems. Native to Asia (China, Japan, Korea), it was introduced to North America at the beginning of the 20^th^ century and then from there to Europe in the 1930’s (Rigling and Prospero, 2018). Additional introductions to Europe as well as to the Caucasus region took place later directly from Asia (Dutech et al., 2012; Prospero and Cleary, 2017). The pathogen spread rapidly in the introduced ranges where it encountered susceptible *Castanea* species (Anagnostakis, 1992) on which it causes lethal bark lesions (so-called cankers). In North America, American chestnut (*C. dentata*) has virtually become extinct because of chestnut blight (Anagnostakis, 1987). By contrast, in Europe the *C. parasitica* epidemic on European chestnut (*Castanea sativa*) showed a milder course due to the appearance and spread in the *C. parasitica* population of a hyperparasitic mycovirus (CHV-1) that acts as biological control agent, and to higher resistance of *C. castanea* compared to *C. dentata* (Rigling and Prospero, 2018). The pathogen can also survive and readily sporulate on fresh dead wood, which is considered to have an important epidemiological role (Prospero et al., 2006; Meyer et al., 2019). Although *C. parasitica* is considered a primary pathogen only on *C. dentata* and *C. sativa* (Anagnostakis, 1987; Braganca et al., 2011), it has also been occasionally detected on various *Quercus* species in different European countries, such as Greece (Tziros et al., 2015), the Czech Republic (Haltofová et al., 2005), Italy (Biraghi, 1950; Dallavalle and Zambonelli, 1999) and Slovakia (Adamčíková et al., 2010).

Recently, two other species of the genus *Cryphonectria* have been officially detected and described, namely *C. naterciae* in Europe (Braganca et al., 2011), and *C. japonica* in Asia (Myburg et al., 2004). *C. naterciae* was isolated in Portugal from bark cankers on *C. sativa* and orange bark lesions on cork oaks (*Quercus suber*). Its closest relatives inferred by partial sequencing of the ITS region are *C. radicalis* and *C. parasitica* (Braganca et al., 2011). *C. japonica* (previously named *C. nitschkei*) was isolated from *Castanea crenata* in Japan (Liu et al., 2003; Gryzenhout et al., 2009) and from *Quercus* spp. in China, on which it causes bark cankers with orange fruiting structures (Jiang et al., 2019). The pathogenicity of *C. naterciae* and *C. japonica* on *C. sativa* and other European species in the family Fagaceae is, however, still unknown.

In this study, we aimed to compare the parasitic and saprotrophic ability of the Asian *C. japonica* with that of an invasive species already established in Europe (*C. parasitica*) and a native European species (*C. naterciae*). As hosts we selected three species of the Fagaceae, i.e. *C. sativa*, the main host of *C. parasitica*, and pedunculate oak (*Quercus robur*) and European beech (*Fagus sylvatica)*, two widespread species in Central Europe. The results of this study will allow a better understanding of the risk posed by *Cryphonectria* species to common European Fagaceae species.

## 2 Materials and Methods

### 2.1 *Cryphonectria* spp. isolates

The study was conducted using three isolates each of *C. parasitica*, *C. naterciae* and *C. japonica* (**Table 1**). Isolates of *C. parasitica* were selected to represent two frequent vegetative compatibility (vc) types (EU-1 and EU-2) and one rare vc type (EU-12) in Switzerland (Prospero and Rigling, 2012). Genotyping at 10 microsatellite loci (Prospero and Rigling, 2012) showed that each of them belonged to a different multilocus genotype (M2372 to CpMG15, M2671 to CpMG37 and M4023 to CpMG21). Isolates of *C. naterciae* and *C. japonica* originated from Portugal and Japan, respectively, and showed no amplification with the *C. parasitica* specific microsatellite assay.

**Table 1:** *Cryphonectria* spp. isolates used in this study

| Isolate name | WSL collection number | Species | Host | Origin | Experiment | Reference |
| --- | --- | --- | --- | --- | --- | --- |
| M2372 | M2372 | *Cryphonectria parasitica* | *Castanea sativa* | Switzerland | Greenhouse/dormant stems | Prospero & Riglig (2012) |
| M2671 | M2671 | *Cryphonectria parasitica* | *Castanea sativa* | Switzerland | Greenhouse/dormant stems | Prospero & Riglig (2012) |
| M4023 | M4023 | *Cryphonectria parasitica* | *Castanea sativa* | Switzerland | Greenhouse/Dormant stems | Prospero & Riglig (2012) |
| OB5-4 | M9250 | *Cryphonectria japonica* | *Castanea crenata* | Japan | Greenhouse/dormant stems | unpublished |
| IF-4 | M9249 | *Cryphonectria japonica* | *Castanea crenata* | Japan | Greenhouse/dormant stems | unpublished |
| E5-5 | M9251 | *Cryphonectria japonica* | *Castanea crenata* | Japan | Greenhouse/dormant stems | unpublished |
| CO679 | M3671 | *Cryphonectria naterciae* | *Castanea sativa* | Portugal | Dormant stems | Bragança et al. (2011) |
| CO607 | M3659 | *Cryphonectria naterciae* | *Quercus suber* | Portugal | Greenhouse/dormant stems | Bragança et al. (2011) |
| CO612 | M3664 | *Cryphonectria naterciae* | *Quercus suber* | Portugal | Greenhouse/dormant stems | Bragança et al. (2011) |

### 2.2 Seedling experiment

Inoculations were performed on two-year-old seedlings of *C. parasitica*, *Q. robur* and *F. sylvatica* obtained from natural populations (i.e. no clonal material). The plants were placed on charts in a chamber of the WSL biosafety greenhouse, which was set at a temperature of 22 C. A random block design was used for the experiment with weekly rotations of the blocks within the chamber to avoid chamber effects. The *Cryphonectria* spp. isolates were grown on Potato Dextrose Agar (PDA; 39g/L, Difco Laboratories, Detroit USA) for 5 days at 25°C in the dark and then inoculated on the stems of the seedlings, as described in Dennert et al. (2019). Briefly, holes of 0.5 cm diameter were made in the bark using a cork borer, and an agar plug of the same size was inserted with the mycelium towards the stem. A total of five seedlings were inoculated with each isolate and 5 seedlings of each tree species with an agar plug as negative control. The inoculation points were sealed with tape and the developing lesions were measured every two weeks for four months. The length and the width were recorded, and lesion area was calculated with the formula for the ellipse area, since lesions had an approximately elliptical shape (Dennert et al., 2019; Meyer et al., 2019). At the end of the experiment, the final lesion area was recorded and used for further analysis. For the seedlings that died during the experiment, the lesion area at the time point of death was used for further analysis. At the end of the experiment or at the time point of seedling death, all lesions were sampled to re-isolate the inoculated *Cryphonectria* species. The outer part of the bark was removed with a sterile scalpel in a sterile bench and small pieces (approx. 4 × 4 mm) of the necrotic tissue were placed on 1.5% water agar with 40 mg/L streptomycin. The plates were incubated at room temperature in the dark and after 5 days, growing colonies were transferred on PDA plates. The identity of the *Cryphonectria* species grown on the PDA plates was assessed visually from the culture morphology using reference cultures for each species.

### 2.3 Dormant stem experiment

Dormant stems of *C. sativa*, *Q. robur* and *F. sylvatica* (3-5 cm diameter) were obtained from Swiss forests. The stems were cut in 50 cm long segments and ends were sealed with paraffin to avoid desiccation. The *Cryphonectria* spp. isolates were cultured and inoculated on the dormant stems with the same method as described for the seedling experiment. To avoid a stem effect on the experiment, on each stem, one randomly selected isolate of each *Cryphonectria* species was inoculated. Five replicate inoculations were performed for each isolate on each tree species. The stems were placed in plastic boxes on plastic supports and the boxes filled with 4L demineralized water to avoid desiccation of the stems (Bryner et al., 2012). The boxes were placed at 25°C in the dark for two weeks. Afterwards, the bark around the inoculation point was removed using a scalpel and the length and width of the lesions that had formed was measured. Since the lesions were approximately elliptical, their area was calculated by using the formula for the ellipse area.

### 2.4 Data analysis

Data were analysed with R version 3.4.1 (R Core Team, 2014). Both in the seedling experiment and in the dormant stem experiment, differences between host-pathogen combinations were analysed with a linear model with tree species and *Cryphonectria* species as factors. The isolates were nested within the *Cryphonectria* species. In the greenhouse experiment, one *C. naterciae* isolate (CO679) had to be excluded from the analyses because it was mixed up with a *C. parasitica* isolate. In the dormant stem experiment, this error was corrected and the measurements for *C. naterciae* CO679 could be included in the analysis.

## 3 Results

### 3.1 Seedling experiment

Mortality was only observed *on* chestnut seedlings inoculated with *C. parasitica* (13 of 15 seedlings died before the end of the experiment). Moreover, this host-pathogen combination resulted in significantly greater lesion areas (average area for single isolates ranged from 26.0 cm^2^ to 34.4 cm^2^) at the end of the experiment than all other combinations (**Figs 1** and **2**). On oak and beech, no mortality was observed and lesion growth was limited, with the smallest lesions measured on beech for all three *Cryphonectria* species (**Fig 2**). Although *C. naterciae* and *C. japonica* produced significantly smaller bark lesions on chestnut than *C. parasitica* (**Fig 1**), they could establish on the seedlings and at the end of the experiment were successfully re-isolated from 80-100% of the lesions (**Fig. 3**). The re-isolations were successful from 20-60% of the lesions on oak for *C. parasitica*, from 40-60% of the lesions for *C. naterciae* and from 0-20% of the lesions for *C. japonica* (**Fig 3**). From beech, the proportion of re-isolations was smaller than those from oak, varying from 0% to 40% of lesions for all three *Cryphonectria* species.

**Figure 1:**
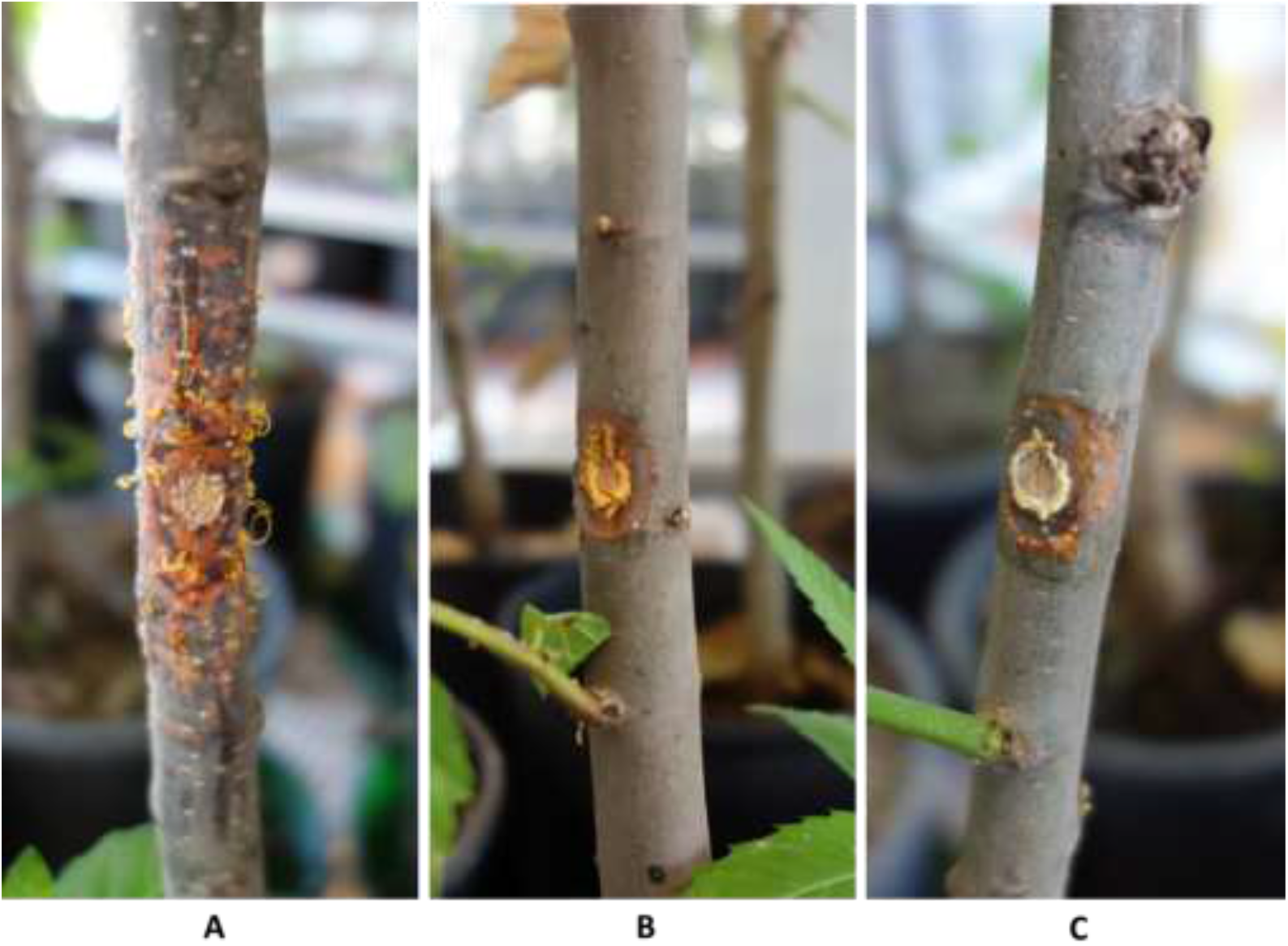
Bark lesions on the stems of two-year-old *Castanea sativa* seedlings caused by *Cryphonectria parasitica* (A), *C. naterciae* (B), and *C. japonica* (C) 1.5 months after inoculation. Abundant asexual sporulation is visible on the seedling inoculated with *C. parasitica* (A). Stems were artificially inoculated with one disk (5 mm diameter) taken from a pure culture of each *Cryphonectria* species grown on PDA.

**Figure 2:**
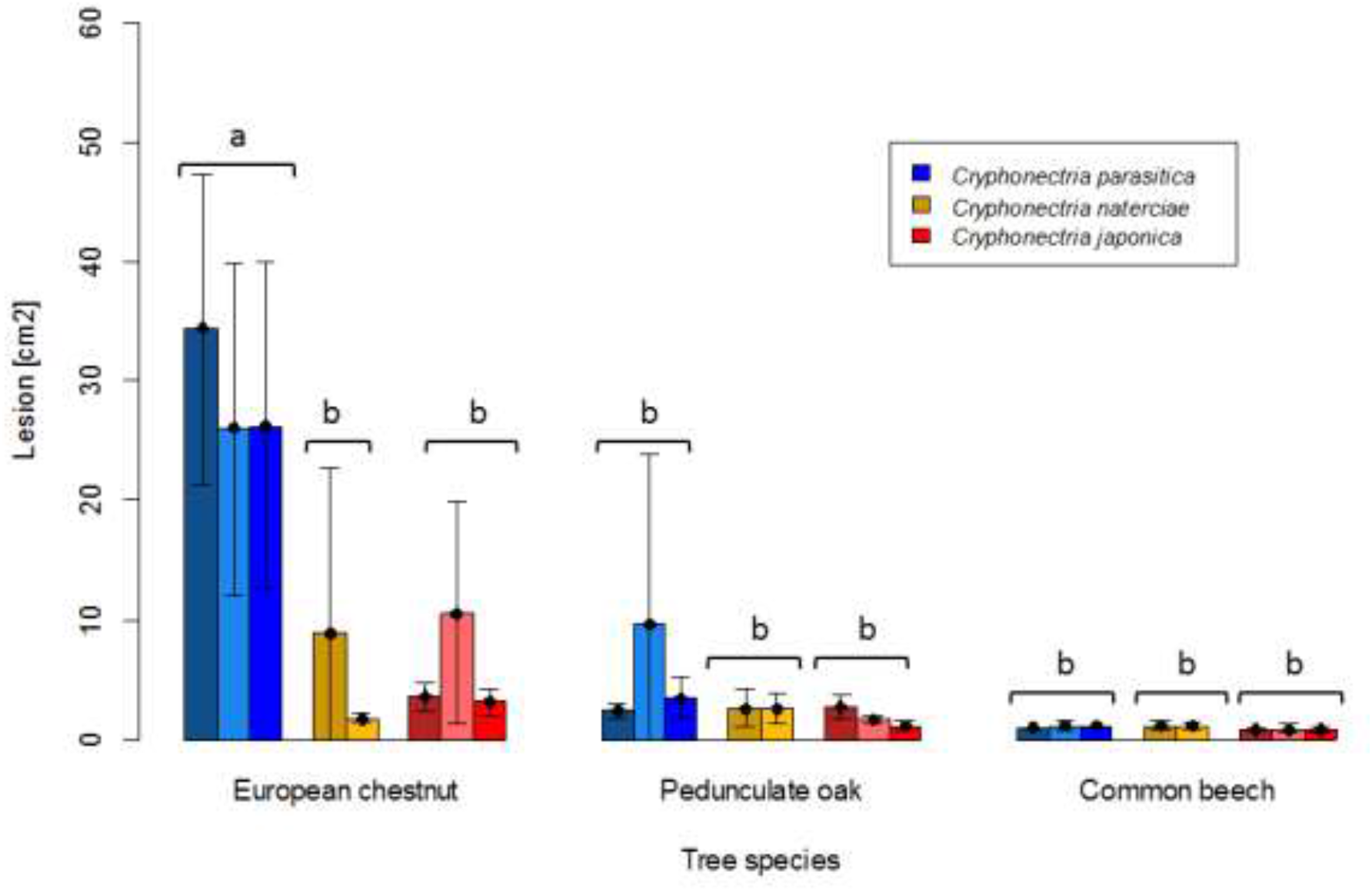
Barplots of lesion sizes caused by *Cryphonectria parasitica* (blue), *C. naterciae* (yellow), and *C. japonica* (red) on the bark of European chestnut (*Castanea sativa*), pedunculate oak (*Quercus robur*) and common beech (*Fagus sylvatica*). Three isolates of *Cryphonectria parasitica* and *C. japonica* and two isolates of *C. naterciae* were tested in the experiment. Each bar shows the average value for one isolate (five seedlings per isolate were inoculated). Black lines show standard deviations. Letters show significant differences between combinations of *Cryphonectria* species and tree species (Linear model with Tukey post-hoc test, p<0.05).

**Figure 3:**
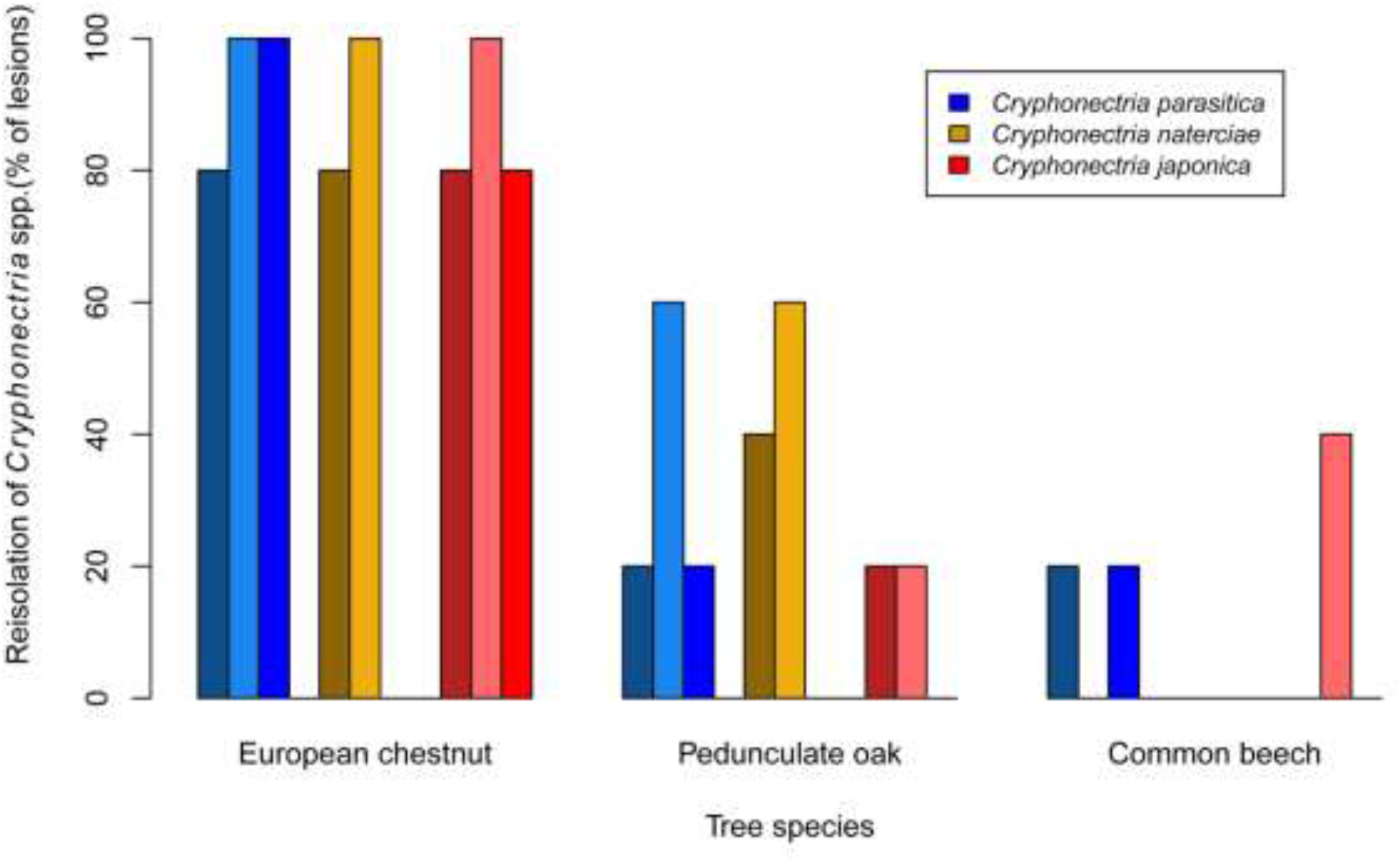
Barplots showing the proportion of bark lesions from which *Cryphonectria parasitica* (blue), *C. naterciae* (yellow), and *C. japonica* (red) could be re-isolated four months after artificial inoculation. Each bar represents one isolate (five seedlings per isolate were used, i.e. the proportion of lesions from which re-isolations were successful is 100% if *Cryphonectria* sp. could be re-isolated from all five lesions).

### 3.2 Dormant stem experiment

In the dormant stem experiment, where the three *Cryphonectria* species were inoculated on chestnut, oak and beech stems to assess saprophytic growth, the results were similar to the seedling experiment for oak and beech. None of the three *Cryphonectria* species could form lesions on these two tree species (**Figs. 4** and **5**). For chestnut, however, the results of the stem experiment differed from those of the seedling experiment, with all three *Cryphonectria* species forming bark lesions on the inoculated stems (**Figs. 4** and **5**). The lesions caused by *C. parasitica* and *C. naterciae* were of similar size (*C. parasitica* average: 47.1 cm^2^, *C. naterciae* average: 52.0 cm^2^), while the lesions caused by *C. japonica* were significantly smaller (average: 22.2 cm^2^), but were nonetheless significantly greater than the lesions induced by all three species on oak and beech (**Fig 5**). Considering the single isolates, in all the three *Cryphonectria* species there seemed to be no clear correlation between the growth on living seedlings and on dormant stems. However, this could not be statistically analysed since the number of isolates per species was too low.

**Figure 4:**
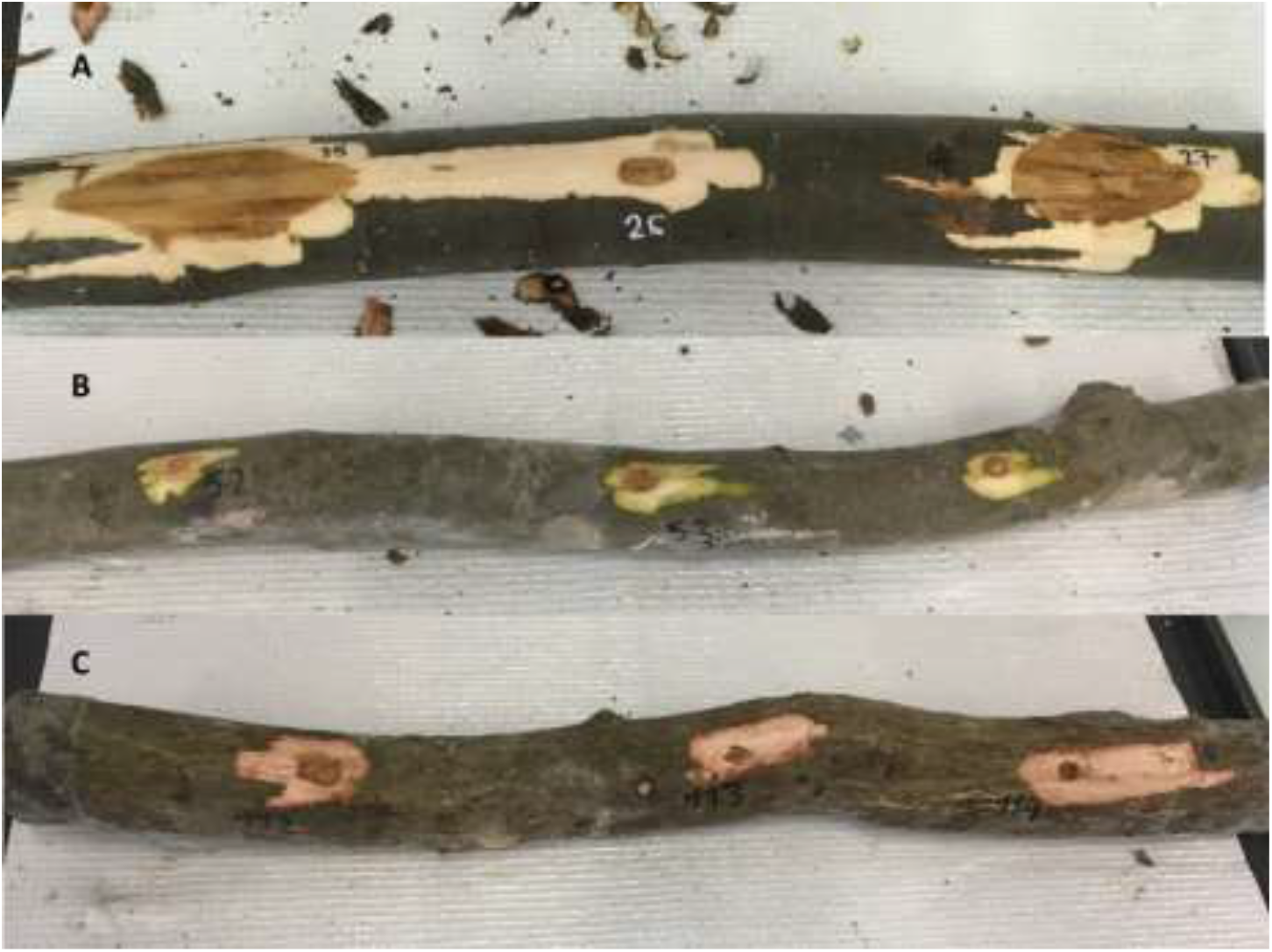
Bark lesions on dormant *Castanea sativa* (A), *Fagus sylvatica* (B) and *Quercus robur* (C) stems obtained after artificial inoculation of *Cryphonectria parasitica*, *C. naterciae* and *C. japonica*. To avoid a stem effect, each stem was inoculated with one randomly selected isolate of each *Cryphonectria* species. For example on stem A, lesion 25 was produced by *C. parasitica*, lesion 26 by *C. japonica*, and lesion 27 by *C. naterciae*.

**Figure 5:**
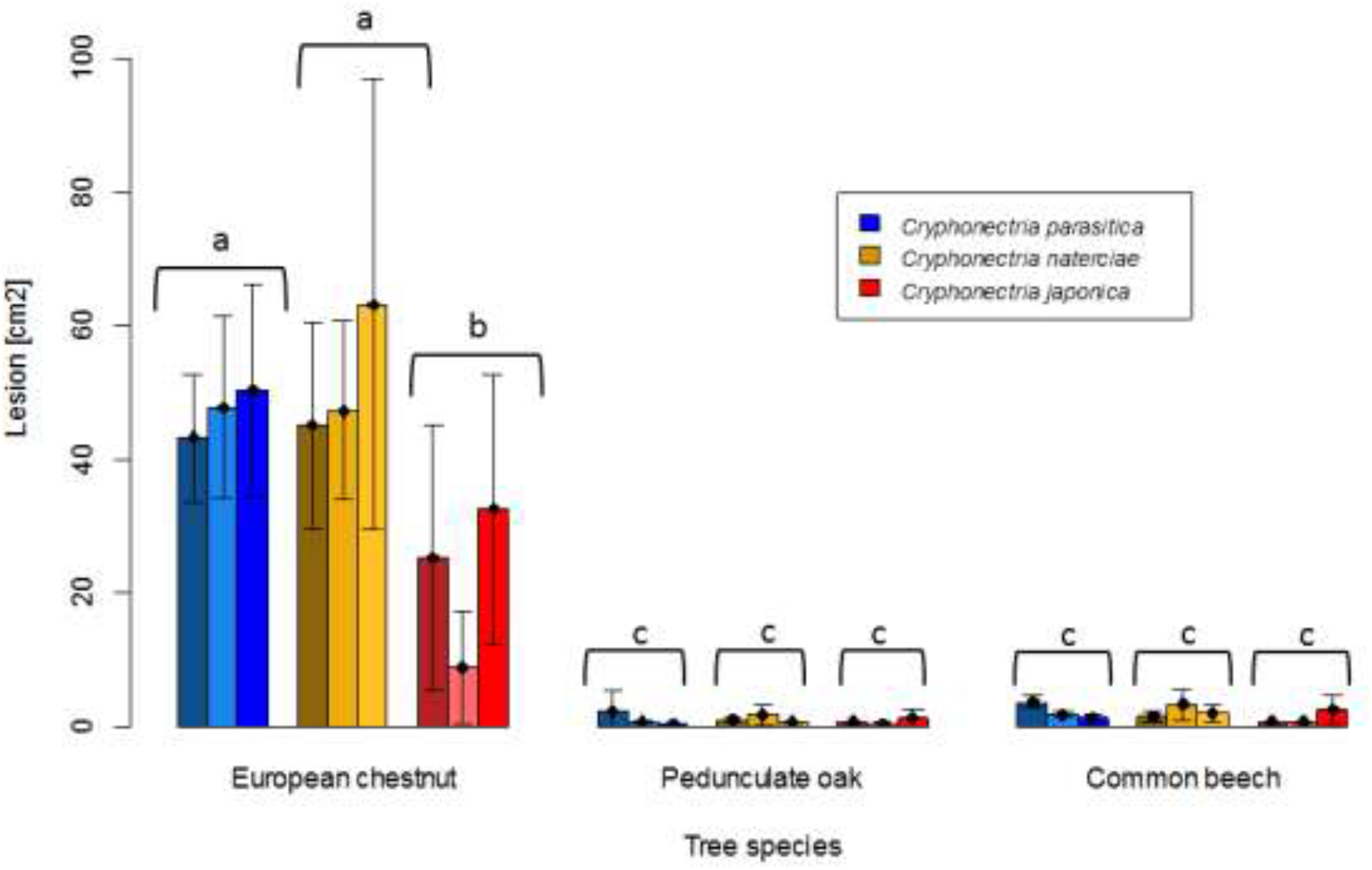
Barplot of lesion sizes obtained on dormant stems of European chestnut (*Castanea sativa)*, pedunculate oak (*Quercus robur*) and common beech (*Fagus sylvatica)* after artificial inoculation of *Cryphonectria parasitica* (blue), *C. naterciae* (yellow) and *C. japonica* (red). Each bar represents the mean lesion size obtained for one isolate (five replicates). Black bars represent standard deviations. Letters show significant differences between combinations of *Cryphonectria* species and tree species (Linear model with Tukey post-hoc test, p<0.05).

## 4 Discussion

To avoid the introduction of new invasive pests, including pathogens, preventive strategies have to be adopted, like border controls and regulated plant trade (Klapwijk et al., 2016). The implementation of such measures, however, implies the identification of potential invasive pests before they become invasive, i.e. before they are moved to a new range. This is particularly challenging because many invasive pests in their native range are harmless endophytes not causing any visible symptoms on their host (e.g. the ash dieback fungus *Hymenoscyphus fraxineus* in Asia; Cleary et al., 2016). Here, we used inoculation experiments to test the pathogenicity of two *Cryphonectria* species closely related to one of the most lethal invasive forest pathogens, namely *Cryphonectria parasitica*. One species (*C. japonica*) is still not present in Europe, whereas the other (*C. naterciae*) seems to be currently restricted to Portugal and some areas in Italy (Braganca et al., 2011; Pinna et al., 2019). Inoculation experiments have previously been used to identify potential new host species of other invasive plant pathogens. For example, Tooley and Kyde (2007) tested the susceptibility to *Phytophthora ramorum* of different potential new host species in the eastern United States. Similarly, Carnegie and Lidbetter (2012), assessed the pathogenicity of an invasive rust species in Australia by inoculating a selection of potential host trees.

Our results show large differences in pathogenicity among the three *Cryphonectria* species included in the study. Differences in lifestyles among congeneric species were previously reported in other fungal genera, including *Alternaria* (Thomma, 2003), *Diaporthe* (Gomes et al., 2013), and *Fusarium* (Kuldau and Yates, 2000; Padhi et al., 2016). Although the Asian species *C. parasitica* is an aggressive pathogen on non-Asian *Castanea* species (Rigling and Prospero, 2018), *C. japonica*, which is also native to Asia, is likely not pathogenic on European chestnut, nor on European beech and pedunculate oak. Indeed, inoculations of *C. japonica* on these three tree species did only result in very small bark lesions. Thus, even if this species would be accidentally introduced into Europe, it would most likely not result in severe damage on the above mentioned tree species. Similarly, *C. naterciae*, a European species present in Portugal and Italy, does not seem to represent a real threat for chestnut, beech and oak, in the case it would be moved outside of its current distribution range. Although *C. naterciae* and *C. japonica* were not pathogenic on seedlings of chestnut and of the other two Fagaceae species, they were able to colonize the bark of freshly cut dormant *C. sativa* stems, and could be successfully re-isolated from the majority of chestnut seedlings in the greenhouse experiment. This suggests that both species may be able to behave as secondary pathogens on *C. sativa* trees already weakened, for example, by abiotic stresses like drought (Desprez-Loustau et al., 2006). Considering the climatic changes predicted for European chestnut growing regions, like an increase of extreme weather events and of drought periods (Giorgi and Lionello, 2008; Mariotti et al., 2015), it would be worth to further investigate the effect of such tree stressors on the pathogenicity of these two *Cryphonectria* species. As observed in *C. parasitica* (Prospero et al., 2006; Meyer et al., 2019), the ability of *C. naterciae* and *C. japonica* to colonize and saprophytically grow on recently dead chestnut stems could contribute in building up a reservoir of inoculum. This could help both species to become successfully established and spread in European forests.

Our study confirms the potential of artificial inoculation experiments to assess the pathogenicity of plant-associated fungi toward selected host species. The tested fungi can include known pathogens, but also saprotrophs or endophytes, which may spread with plants for planting and represent a threat for non-native hosts. This approach is particularly helpful in combination with sentinel plantings, where host species are grown outside their natural distribution range to detect potential new pathogens (Morales-Rodríguez et al., 2019). Once such pathogens have been identified, their host range can be determined by conducting inoculation experiments with living plants in the greenhouse. However, invasiveness of a fungal pathogen (i.e. the ability to arrive, spread beyond its introduction site and become established in a new location; Desprez-Loustau et al., 2007) is not only a function of its pathogenicity. Other phenotypic traits, such as sporulation and saprotrophic ability may play an important role. Moreover, environmental conditions strongly influence plant diseases, therefore climate changes can also be considered drivers of disease outbreaks (Sturrock et al., 2011). Thus, to obtain a full view it is necessary to experimentally assess as many traits as possible. In our specific case, artificial inoculations showed that an accidental introduction to Europe of *C. japonica* or the spread within Europe of *C. naterciae* would most likely have much less devastating consequences than the invasion by *C. parasitica*. Nonetheless, the ability of both species to saprotrophically colonize fresh chestnut wood may suggest that they could become established in chestnut forests and eventually infect weakened chestnut trees or other hosts not tested in this study.

## 5 Conflict of Interest

The authors declare that the research was conducted in the absence of any commercial or financial relationships that could be construed as a potential conflict of interest.

## 6 Author Contributions

FD collected and analysed the data of the dormant stem experiment and wrote the first version of the manuscript, DR planned the seedling experiment together with JBM and SP and performed the re-isolations from the inoculated seedlings, JBM performed the seedling experiment with SP, CS analysed the data of the seedling experiment, EA performed the dormant stem experiment, SP planned the experiments and supervised the study. All authors contributed to the final version of the manuscript.

## 7 Funding

This project was funded by the Swiss Federal Institute for Forest, Snow and Landscape Research (WSL).

## 8 Acknowledgments

We thank Helena Braganca for providing the *C. naterciae* isolates and Michael Milgroom for providing the *C. japonica* isolates used in this study. We also thank Hélène Blauenstein for technical support in the WSL biosafety facility.

